# KIT is required for fetal liver erythropoiesis but dispensable for angiogenesis

**DOI:** 10.1101/2021.01.15.426801

**Authors:** Alessandro Fantin, Alice Plein, Carlotta Tacconi, Emanuela Villa, Elena Ceccacci, Laura Denti, Christiana Ruhrberg

## Abstract

Blood vessels are fundamental to sustain organ growth and tissue metabolism. In the mouse embryo, endothelial cell (EC) progenitors almost concomitantly give rise to the first blood vessels in the yolk sac and the large vessels of the embryo proper. Thereafter, the vascular network expands by angiogenesis to vascularize developing organs such as the brain. Although the first blood cells form in the yolk sac before blood vessels have assembled, consecutive waves of hematopoietic progenitors subsequently bud from hemogenic endothelium located within the wall of yolk sac and large intraembryonic vessels in a process termed endothelial to hematopoietic transition (endoHT). The receptor tyrosine kinase KIT is required for late embryonic erythropoiesis, but KIT is also expressed earlier in the hemogenic endothelium, in hematopoietic progenitors that arise via endoHT from hemogenic endothelium and non-hemogenic ECs, such as in the brain. However, it remains unclear whether KIT has essential roles in early hematopoiesis or even blood vessel growth. Here, we have combined transcriptomic analysis to delineate *Kit* expression with the analysis of knockout mice to show that KIT is expressed during but dispensable for yolk sac endoHT or brain angiogenesis but required for transient definitive erythropoiesis in the fetal liver.

## Introduction

In all vertebrates, endothelial cells (ECs) give rise to the blood vasculature, which supplies nutrients and oxygen whilst removing waste materials. ECs differentiate from mesenchymal precursors termed angioblasts, which in mice appear in the extra-embryonic yolk sac on embryonic day (E) 7.0 and almost concomitantly arise in the lateral plate mesoderm of the embryo proper (Potente et al., 2011). These early ECs condense into the yolk sac vasculature and dorsal aortae in a process termed vasculogenesis and thereafter migrate and proliferate to expand the vasculature by sprouting angiogenesis (Potente et al., 2011). In close spatiotemporal proximity to blood vessel formation, several consecutive waves of hematopoietic progenitors arise and contribute blood and immune cells to the growing vertebrate embryo (Hoeffel and Ginhoux, 2018).

The first, primitive hematopoietic precursors originate in the yolk sac blood islands between E7.0 and E8.25 in the mouse and differentiate into embryonic erythrocytes and yolk sac macrophages (Ginhoux and Guilliams, 2016; Hoeffel and Ginhoux, 2018). The onset of blood flow allows yolk sac macrophages to colonize the embryo and differentiate into tissue-resident macrophages. The transient definitive wave of hematopoietic precursors arises when a subset of ECs in the yolk sac specializes into hemogenic endothelium to undergo an endothelial-to-hematopoietic-transition (endoHT) between E8.5 and E9.5 in the mouse (Hoeffel and Ginhoux, 2018). This process generates erythromyeloid progenitors (EMPs), which leave the yolk sac after the onset of blood flow and colonize the liver (Frame et al., 2013; Hoeffel et al., 2015; Hoeffel and Ginhoux, 2018; Lux et al., 2008; McGrath et al., 2015). In the liver, EMPs give rise to both transient definitive erythrocytes and monocyte precursors for tissue macrophages. It is thought that the liver EMP-derived macrophages replace the initial pool of yolk sac-derived macrophages in all organs, with the exception of the tissue-resident macrophages of the central nervous system, termed microglia (Hashimoto et al., 2013; Hoeffel et al., 2015). EMPs also contribute ECs to organ vasculature (Plein et al., 2018), although different genetic tools to lineage trace EMP-derived cells have yielded conflicting results as to the prevalence of EMP-derived ECs (Feng et al., 2020). Finally, the definitive wave of hematopoietic precursors emerges via endoHT in the aorta-gonad-mesonephros (AGM) region (Sanchez et al., 1996; Yokomizo and Dzierzak, 2010) as a continuum of pro-, pre-and definitive hematopoietic stem cells (HSCs) that seed the liver from E10.5 in the mouse before colonizing the bone marrow after birth (Azzoni et al., 2018; Rybtsov et al., 2014; Rybtsov et al., 2011; Taoudi et al., 2008).

Hemogenic ECs are induced by retinoic acid signaling, which upregulates the expression of the receptor tyrosine kinase KIT (Dejana et al., 2017), whereby KIT cell surface expression is often used as a distinguishing feature from non-blood forming ECs (Goldie et al., 2008). Moreover, KIT is also used as a key marker for the progeny of hemogenic endothelium, including EMPs (Gomez Perdiguero et al., 2015; Hoeffel et al., 2015; McGrath et al., 2015) and HSCs (Gomez Perdiguero et al., 2015; Sanchez et al., 1996). Genetic defects that disrupt KIT signaling reduce the number of late embryonic HSCs (Ikuta and Weissman, 1992) and cause severe anemia and thus late embryonic or perinatal lethality (Bernex et al., 1996; Broudy, 1997; Ding et al., 2012). The anemic phenotype was ascribed to an erythroid differentiation block in the fetal liver after E13.5 (Broudy, 1997; Chui et al., 1978; Chui and Loyer, 1975; Chui and Russell, 1974). Subsequent studies with function-blocking antibodies further suggested that hematopoietic waves originated before E12.5 are less dependent on KIT than later embryonic waves (Ogawa et al., 1993). However, it has not been directly addressed whether KIT is required for endoHT and, specifically, EMP formation and function. KIT has also been implicated in EC migration, proliferation and tube formation through experimentation *in vitro* (Kim et al., 2011; Kim et al., 2019; Matsui et al., 2004), and decreased KIT expression reduces angiogenesis in mouse models of ocular pathology (Kim et al., 2019) and tumor growth (Fang et al., 2012). Moreover, it has recently been shown that KIT is expressed in brain endothelium between E9.5 and E11.5 (Feng et al., 2020), corresponding to the time when the brain is first vascularized by sprouting angiogenesis (Fantin et al., 2013b; Tata et al., 2015). However, it has not yet been examined whether KIT is required for embryonic blood vessel growth.

Here, we have performed single cell expression analyses in mouse embryos to identify an organ-specific cellular profile of *Kit* expression during embryogenesis. Our functional studies further show that the hemato-vascular requirement for KIT does not include yolk sac hematopoiesis or brain angiogenesis but extends to the EMP-dependent transient-definitive erythropoiesis that takes place in the fetal liver. KIT is therefore required for erythropoiesis earlier than previously reported.

## Results

### KIT marks the hemogenic EC and EMP surface but is dispensable for endoHT in the yolk sac

Wholemount immunostaining of E9.5 yolk sacs localized KIT to small clusters of cells within the CDH5^+^ KDR^+^ endothelium that appeared rounder and smaller than neighboring ECs, consistent with imminent budding into the vascular lumen (**Fig. 1a**). These observations support that KIT is expressed by hemogenic ECs undergoing endoHT and that KIT immunostaining distinguishes hemogenic and non-blood forming ECs. We further found that CDH5 was concentrated at adherens junctions at cell-cell contacts between KIT^−^ ECs, whilst KIT^+^ ECs had more intracellular CDH5 staining than neighboring ECs. This finding suggests that CDH5 internalization precedes EMP budding.

**Figure 1.**
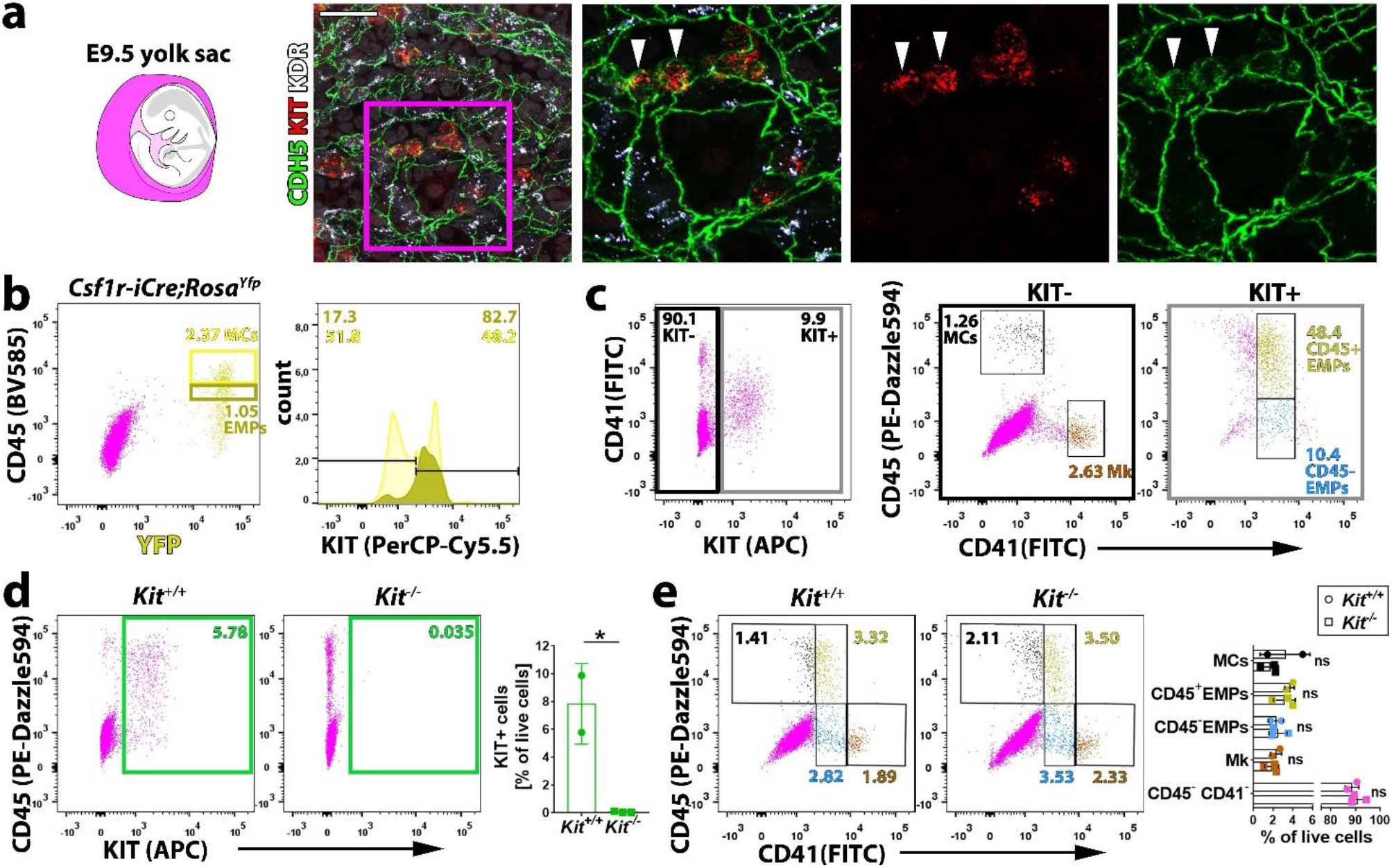
KIT is expressed by hemogenic ECs and EMPs in the yolk sac but is dispensable for yolk sac hematopoiesis. (**a**) Wholemount labelling of E9.5 mouse yolk sacs with the indicated markers (n = 2); arrowheads indicate KIT^+^ budding EMPs; scale bar: 50 μm. (**b**) Flow cytometry analysis of single living cells from E9.5 *Csf1r-iCre;Rosa^Yfp^* mouse yolk sacs with the indicated markers (n = 1 of 4 pooled yolk sacs). The dark and bright yellow insets in the scatter plot indicate putative CD45^low^ YFP^+^ EMPs and CD45^high^ YFP^+^ myeloid cells (MCs) and the gates used to generate the respective histograms. (**c**) Flow cytometry analysis of single living cells from E9.5 mouse yolk sacs with the indicated markers (n = 2). The black and grey insets in the scatter plot on the left hand side indicate the gates used to generate the scatter plots in the center and on the right hand side, respectively; Mk, megakaryoblasts; EMP, erythro-myeloid progenitors; MCs, myeloid cells. (**d,e**) Flow cytometry analysis of single living cells from E9.5 *Kit^+/+^* (n = 2) and *Kit^−/−^* (n = 3) littermate mouse yolk sacs with the indicated markers. (**d**) Quantification of KIT^+^ cells, as shown in the green insets, as well as cells defined by CD45 or CD41 expression levels (**e**), as shown in the color-coded scatter plots. All bar graph data are shown as mean ± SD; each data point represents the value from one embryo; ns, non-significant (2way ANOVA followed by Sidak’s multiple comparisons test), *p<0.05 (unpaired Student’s t-test).

To identify EMPs with flow cytometry, KIT surface expression has previously been combined with low levels of CD45 (CD45^low^) and *Csf1r-iCre*-mediated lineage tracing using recombination reporters such as *Rosa^Yfp^* (Gomez Perdiguero et al., 2015; Hoeffel et al., 2015) or with CD41 co-expression (Frame et al., 2016; McGrath et al., 2015). We found that more than 80% of CD45^low^ YFP^+^ cells in E9.5 *Csf1r-iCre;Rosa^Yfp^* yolk sacs were also KIT^+^ (**Fig. 1b**). Analysis of KIT surface expression with both CD41 and CD45 identified a KIT^−^ cell population (**Fig. 1c**) comprised of CD45^+^ CD41^−^ cells that likely correspond to myeloid cells (Frame et al., 2016; McGrath et al., 2015) and CD45^−^ CD41^+^ cells that likely corresponds to megakaryoblasts (Cortegano et al., 2019; Frame et al., 2016; McGrath et al., 2015). By contrast, the KIT^+^ population (**Fig. 1c**) included CD41^low^ cells, which were mostly CD45^+^. This cell population likely contains progenitors with definitive erythroid, myeloid and lymphoid potential, including the previously described *Csf1r*-lineage traced CD45^low^ KIT^+^ EMPs (Gomez Perdiguero et al., 2015; Hoeffel et al., 2015) and CD45^+^ CD41^+^ KIT^+^ EMPs (Frame et al., 2016; McGrath et al., 2015), as well as lymphomyeloid progenitors (Boiers et al., 2013). The smaller proportion of KIT^+^ CD41^low^ progenitors that did not express CD45 (**Fig. 1c**) may include EMPs that are CD45^−^ (McGrath et al., 2015), possibly because they are less mature, as well as other KIT^+^ CD41^+^ CD45^−^ hematopoietic precursors that have recently been described (Yamane, 2018).

To understand whether KIT promotes EMP formation or affects the differentiation potential of EMPs, we next performed flow cytometry analysis of yolk sacs from mice lacking KIT. For this, we used mice homozygous for the *Kit^CreERT2^* knock-in allele, in which the *Cre* recombinase gene is inserted into the endogenous *Kit* locus (Klein et al., 2013) to generate a true *Kit* null allele (Heger et al., 2014). These mice are hereafter referred to as *Kit^-/-^* mutants. Notably, KIT loss in E9.5 yolk sacs (**Fig. 1d**) did not significantly alter the proportion of the CD45^−^ CD41^−^ population that includes erythroblasts (Frame et al., 2016), the CD45^−^ CD41^+^ population that includes megakaryoblasts, the CD41^low^ progenitor population that includes both CD45^+^ and CD45^−^ EMPs or the CD45^+^ CD41^−^ population corresponding to differentiating myeloid cells (MCs) (**Fig. 1e**). These observations suggest that KIT is dispensable for hematopoiesis and therefore also endoHT in the yolk sac.

### KIT is expressed in embryonic brain ECs, but not required for brain angiogenesis

*Kit* transcripts and KIT protein were recently described to be present in embryonic brain endothelium (Feng et al., 2020). To corroborate this finding and place it into context of brain vascular development, we analyzed *Kit* expression in publicly available single cell RNA sequencing (scRNA-seq) datasets from the E11.5 and E12.5 mouse midbrain (GSE76381). For both time points, we performed graph-based clustering. Our annotation of the E11.5 dataset identified ECs (*Pecam1^+^*) alongside pericytes (*Pdgfrb^+^*), a neural population comprised of neural progenitors (*Cenpf^+^*), neuroblasts (*Onecut2^+^*) and motor neurons (*Isl1^+^*), as well as microglia (*Csf1r^+^*) (**Fig. 2a**). Notably, the transcriptomes of these cell populations all formed discrete clusters when Uniform Manifold Approximation and Projection (UMAP) was applied to reduce dimensionality (**Fig. 2b**). Consistent with active angiogenesis in the E11.5 brain (Fantin et al., 2010; Fantin et al., 2013b; Tata et al., 2015), the EC cluster included cells with transcripts for the endothelial tip cell markers *Dll4* and *Apln* and the proliferation markers *Pcna* and *Mki67* (**Fig. 2c**). In contrast, *Runx1*, a marker of hemogenic ECs of the yolk sac and dorsal aorta (Chen et al., 2009; Frame et al., 2016; Yzaguirre et al., 2017), was not obviously expressed in E11.5 brain ECs (**Fig. 2c**). Although *Runx1* expression was not detectable, the majority of E11.5 brain ECs nevertheless contained *Kit* transcripts (**Fig. 2c,d**). Similar results were obtained at E12.5 (**Fig. 2e-h**), except that a microglia signature was not retrieved, possibly because this data set was obtained from fewer cells. Lack of *Runx1* expression at E11.5 and E12.5 suggests that brain vasculature is not hemogenic, despite recent reports hypothesizing otherwise (Gama Sosa et al., 2020; Li et al., 2012). *Kit* expression in brain EC in the absence of *Runx1* expression at both ages instead raises the possibility that KIT plays a role in ECs during brain vascularization.

**Figure 2.**
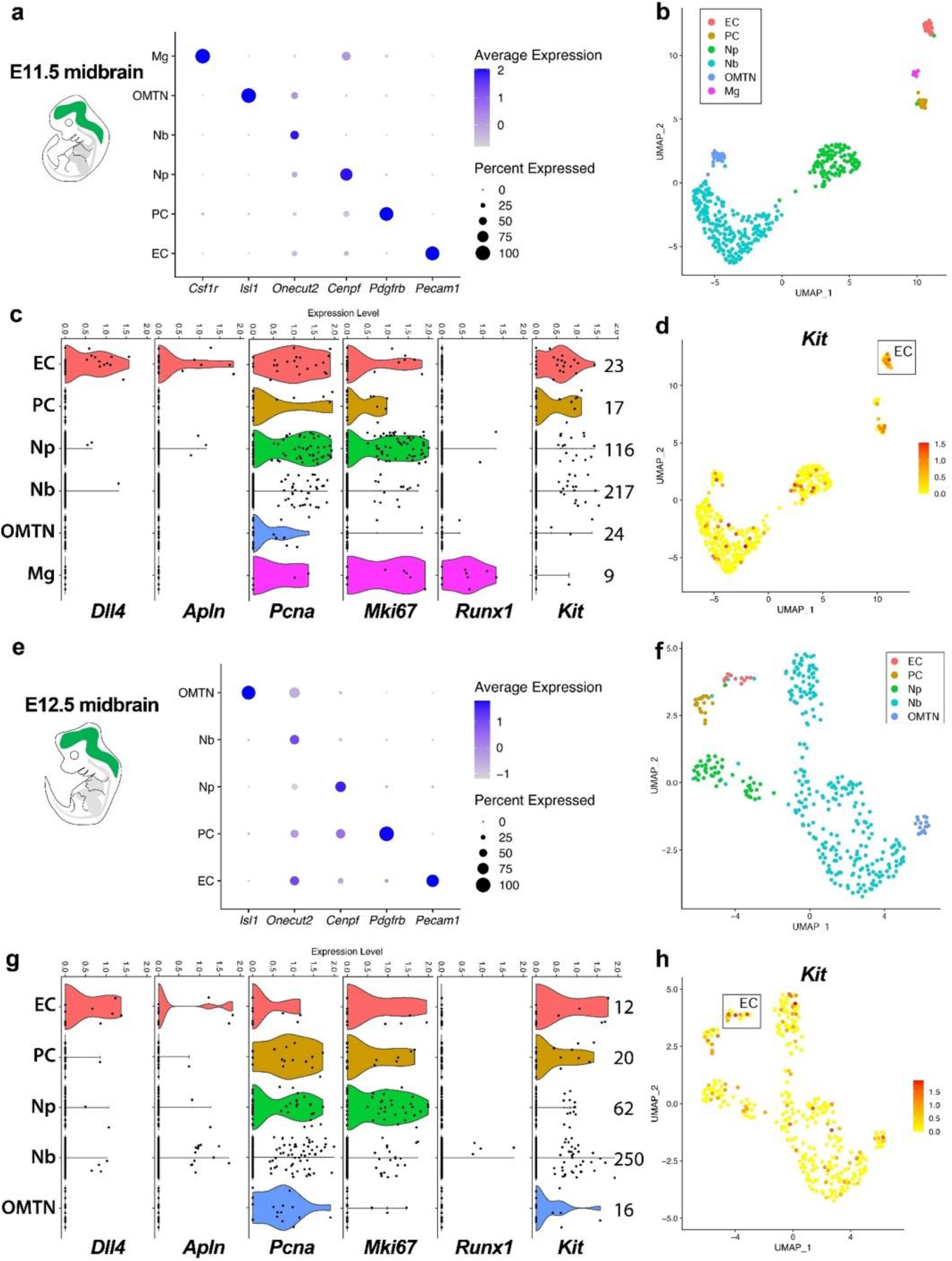
*Kit* expression in major cell types of the embryonic brain. scRNA-seq analysis of E11.5 and E12.5 mouse midbrain. (**a,e**) Expression of the indicated marker genes for the indicated cell types, shown as bubble plots; the color intensity represents the average transcript level, whilst the size of each dot corresponds to the percentage of cells in which that marker was detected. (**b-d,f-h**) UMAP plots visualize clusters of distinct cell types (**b,f**) and *Kit* transcript levels (**d,h**) in each cell cluster. Violin plots (**c,g**) illustrate transcript levels for the indicated genes, whereby the number of cells in each cluster is indicated to the right of the violin plots. Mg, microglia; PC, pericytes; Np, neural progenitors; Nb, neuroblasts; OMTN, oculomotor/trochlear nucleus.

To determine whether KIT is required for brain angiogenesis, we used the hindbrain as an established angiogenesis model (Fantin et al., 2013b). Fluorescent wholemount staining for the vascular endothelial marker isolectin B4 (IB4) revealed similar vascular complexity in *Kit^+/+^* and *Kit^−/−^* hindbrains (**Fig. 3a,b**). We also compared the contribution of *Csf1r-iCre* lineage ECs to *Kit^+/+^* and *Kit^−/−^* hindbrains, because we have recently found that KIT^+^ *Csf1r*^+^, EMP-like vascular progenitors contribute to brain angiogenesis (Plein et al., 2018). However, the proportion of tdTomato^+^ ECs in *Csf1r-iCre;Rosa^tdTom^* hindbrain vasculature was similar in *Kit^+/+^* and *Kit^−/−^* hindbrains (**Fig. 3a,c**). Moreover, the number of microglia was not different between *Kit^+/+^* and *Kit^−/−^* hindbrains (**Fig. 3a,d**). As microglia arise from yolk sac-born macrophages before liver colonization by EMPs, this observation agrees with the idea that KIT is dispensable for yolk sac hematopoiesis (**Fig. 1**).

**Figure 3.**
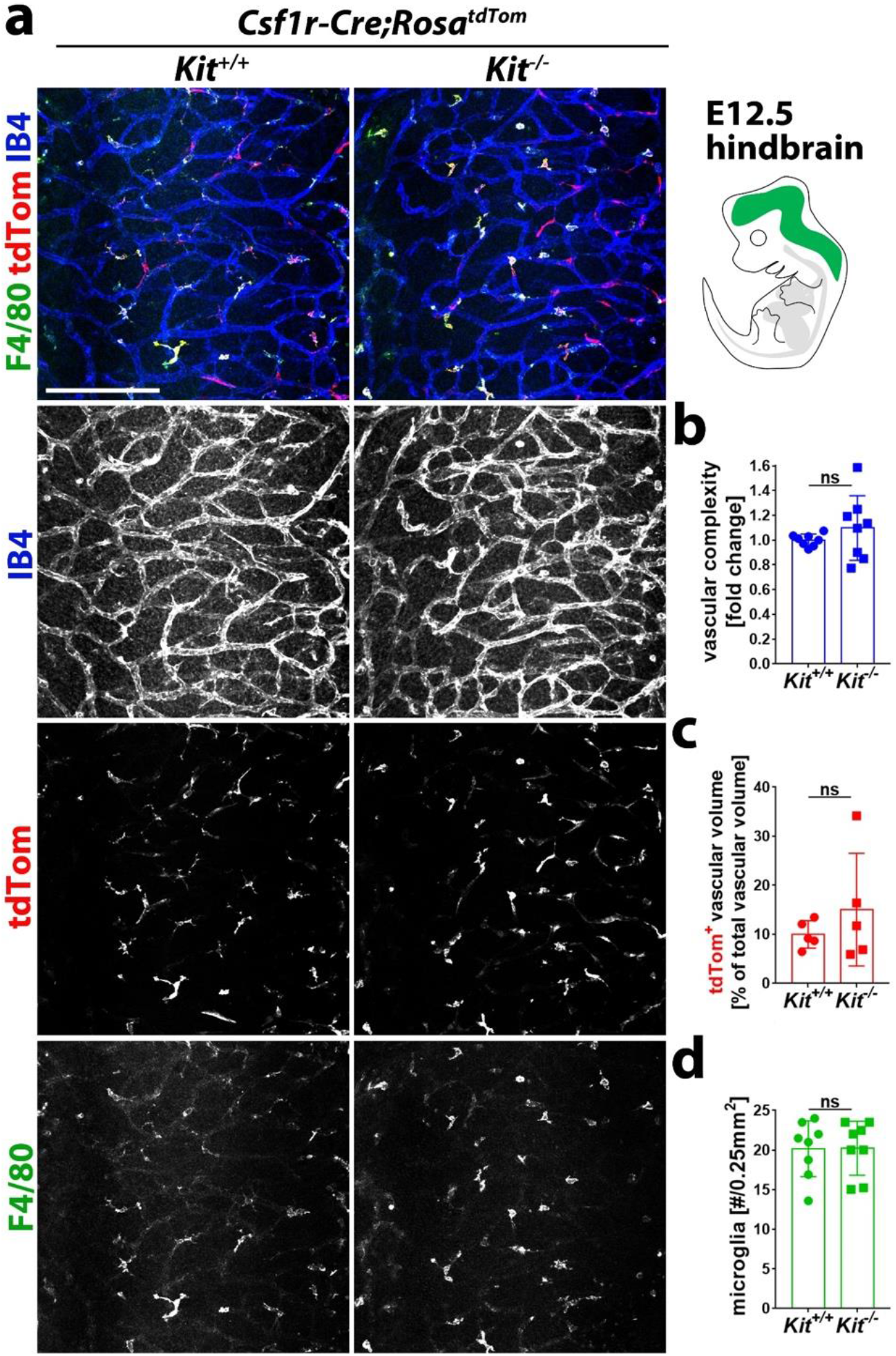
KIT is dispensable for brain angiogenesis, *Csf1r* lineage EC contribution and microglia development. E12.5 mouse hindbrains from *Kit^+/+^* versus *Kit^−/−^* littermates with the *Csf1r-iCre;Rosa^tdTom^* reporter were stained with the indicated markers and imaged by confocal microscopy (**a**) before quantification of the indicated vascular parameters (**b-d**); n = 8 wildtype and mutant hindbrains each, except for tdTomato^+^ samples (n = 5 each). Single channel images are also shown in grey scale. Scale bar: 200 μm. (**b**) Vascular complexity is shown as fold change of vascular branchpoints in mutants relative to wild type. (**c**) tdTomato^+^ EC volume relative to the total IB4^+^ EC volume. (**d**) Number of F4/80^+^ microglia in each 0.25 mm^2^ area. Bar graph data are shown as mean ± SD; each data point represents the value from one embryo; ns, non-significant (unpaired Student’s t-test); tdTom, tdTomato.

### *Kit* is required for transient-definitive fetal liver erythropoiesis

Starting from E10.5 onwards, EMPs leave the yolk sac and colonize the fetal liver (Frame et al., 2013; Hoeffel et al., 2015; Hoeffel and Ginhoux, 2018; Lux et al., 2008; McGrath et al., 2015). To identify EMPs in E12.5 *Csf1r-iCre;Rosa^tdTom^* liver, we classified them as CD45^+^ tdTomato^+^ cells lacking the differentiated myeloid marker CD11b and found that ~90% of these cells were KIT^+^ (**Fig. 4a,b**). In contrast, CD45^+^ tdTomato^+^ CD11b^+^ cells mostly lacked KIT (**Fig. 4a,b**).

**Figure 4.**
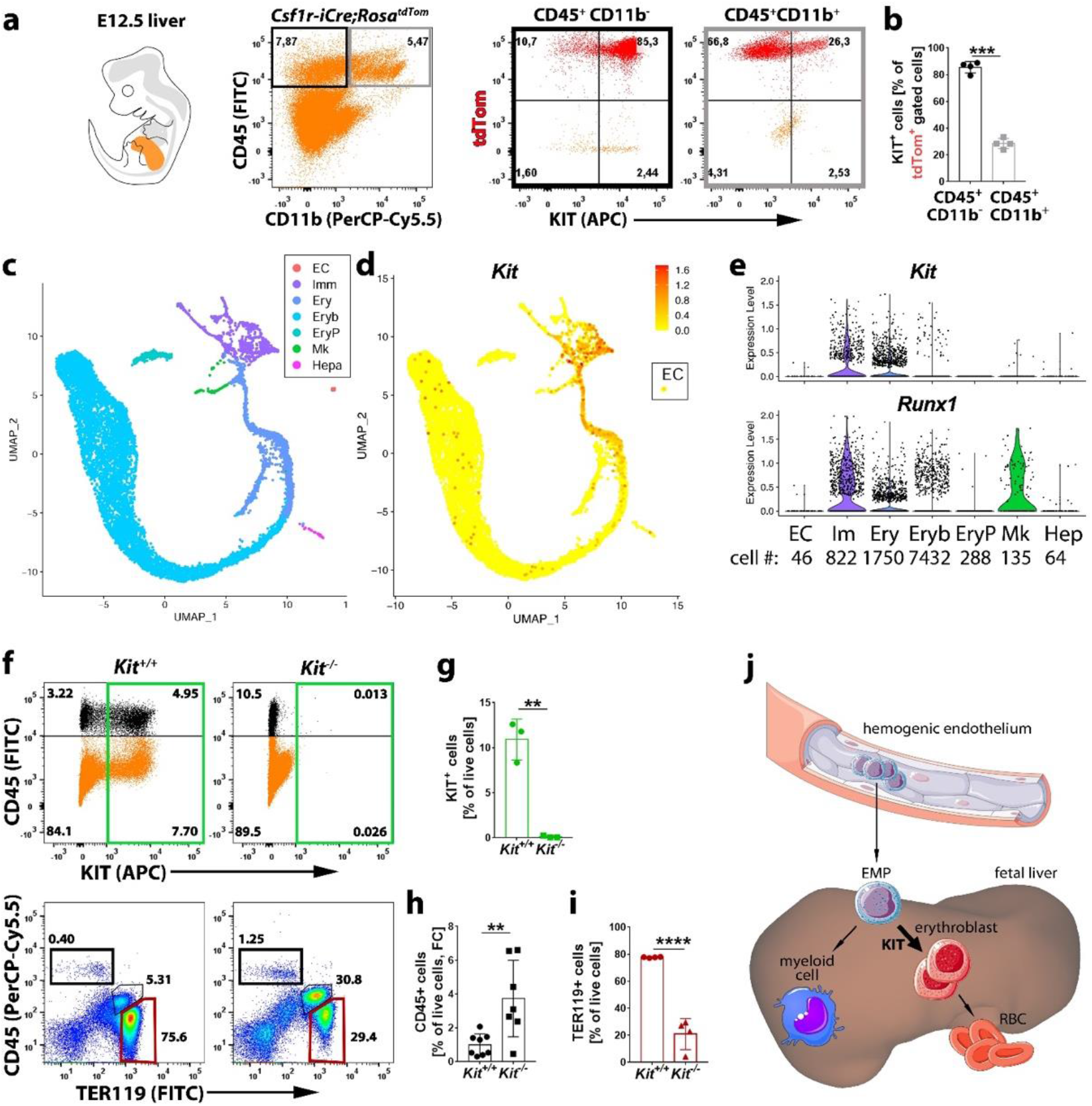
*Kit* expression profiling and requirement during fetal liver hematopoiesis. (**a**) Flow cytometry analysis of single living cells from E12.5 *Csf1r-iCre;Rosa^tdTom^* fetal mouse livers with the indicated markers. The black and grey insets in the left scatter plot indicate the gates used to generate the scatter plots of CD45^+^ cells lacking or expressing CD11b (right). (**b**) Quantification of KIT^+^ tdTomato^+^ cells in the CD45^+^ CD11b^−^ and CD45^+^ CD11b^+^ populations (n = 4 livers). Data are shown as mean ± SD; each data point represents the value from one embryo; ***p<0.001 (paired Student’s t-test). (**c-e**) scRNA-seq analysis of E12.5 mouse liver. UMAP plots were used to visualize clusters of distinct cell types (**c**) and *Kit* transcript levels in each cell (**d**). The EC cluster in (**d**) is indicated with a box. (**e**) Violin plots illustrate *Kit* and *Runx1* single cell transcript levels in each cluster; the number of cells in each cluster is indicated below the violin plots. (**f**) Flow cytometry analysis of single living cells from E12.5 *Kit^−/−^* and *Kit^+/+^* littermate mouse livers with the indicated markers. (**g-i**) Quantification of KIT^+^ (**g**; n = 3 each), CD45^+^ (**h**; n = 8 wild types and n = 7 mutants) and TER119^+^ (**i**; n = 4 each) cells in the gate indicated by the color-coded insets. Data are shown as mean ± SD; each data point represents the value from one embryo; ns, non-significant, **p<0.01, ****p<0.0001 (unpaired Student’s t-test). (**j**) Working model showing how KIT promotes the differentiation of yolk sac hemogenic endothelium-derived EMPs towards the erythroid lineage at the expense of the myeloid lineage.

To define the KIT^+^ cell populations in the fetal liver more clearly, we analyzed the E12.5 liver transcriptome by scRNA-seq (manuscript in preparation). Consistent with the liver being the major hematopoietic organ at E12.5, our annotation identified several hematopoietic cell types in a UMAP continuum that included megakaryocytes (*Pf4*^+^), erythroblasts (*Klf1*^+^ *Rhd*^+^), erythroid-committed progenitors or burst-forming unit erythroid (BFU-E) cells (*Klf1*^+^ *Rhd*^−^) and a cluster of immune cells (*Ptprc*^+^) that included hematopoietic progenitors such as EMPs as well as differentiated myeloid cells (**Fig. 4c** and manuscript in preparation). UMAP cluster continuity suggests that these cells types are likely derived from a shared hematopoietic progenitor lineage in the immune cell cluster. Moreover, we identified distinct clusters of yolk sac-derived primitive orthochromatic erythroblasts (*Hba-x*^+^) as well as hepatoblasts (*Alb*^+^) and ECs (*Cldn5*^+^) (**Fig. 4c**).

We next analyzed which liver cell types expressed *Kit*. Hepatoblasts and fetal liver ECs did not contain *Kit* transcripts at E12.5 (**Fig. 4d,e**). Moreover, fetal liver ECs lacked transcripts for the hemogenic endothelium marker *Runx1* (**Fig. 4e**). These findings agree with the concept that the fetal liver lacks hemogenic endothelium, but harbors recruited hematopoietic progenitors (Frame et al., 2013; Hoeffel et al., 2015; Hoeffel and Ginhoux, 2018; Lux et al., 2008; McGrath et al., 2015). *Kit* transcripts were also not detected in primitive erythroblasts or in differentiating megakaryocytes (**Fig. 4d,e**). However, *Kit* transcripts were abundant in erythroid-committed progenitors and in the immune cell cluster that includes differentiated myeloid cells and EMPs (**Fig. 4d,e**). Notably, the number of *Kit*^+^ cells gradually decreased in erythroid-committed progenitors while they differentiated towards an erythroblast phenotype and appeared dramatically reduced in the erythroblast cluster (**Fig. 4d,e**). This expression pattern is compatible with a selective role for *Kit* in the hematopoietic progenitors that underpin fetal liver erythropoiesis, including the erythroid-committed progeny of EMPs.

Flow cytometry analysis of E12.5 livers confirmed that KIT is expressed in both the CD45^+^ population that includes differentiated myeloid cells and EMPs as well as in the CD45^−^ population that includes TER119^+^ erythroid cells (**Fig. 4f**). KIT loss significantly decreased the proportion of TER119^+^ cells while concomitantly increasing the percentage of CD45^+^ cells (**Fig. 4g-i**). The imbalanced production of TER119^+^ erythroid versus CD45^+^ immune/myeloid cells suggests that KIT plays a specific role in fate decisions made by EMPs as they differentiate in the fetal liver and therefore shows that KIT is required for fetal liver erythropoiesis (**Fig. 4j**).

## Discussion

KIT has been shown to be required for hemogenic EC specification and function *ex vivo* (Marcelo et al., 2013). Indeed, our analysis showed KIT localization to a subset of yolk sac endothelial cells which appear to give rise to budding hematopoietic progenitors at E9.5 (**Fig. 1**). The budding cells are thought to be mostly EMPs, a recently defined wave of transient definitive hematopoietic progenitors in E8.5 and E9.5 yolk sacs that exhibit erythroid and broad myeloid, but not lymphoid potential (Frame et al., 2013; Ginhoux and Guilliams, 2016). Unexpectedly, however, our flow cytometry analysis of KIT null mutants showed that KIT is dispensable for the generation of CD41^low^ progenitors, which include both CD45^+^ and CD45^−^ KIT^+^ EMPs; further, we found that KIT is dispensable for the immediate differentiation steps of EMPs towards the myeloid, erythroid and megakaryocyte lineages at E9.5 (**Fig. 1**). These findings, in turn, suggest that hemogenic endothelium can form even in the absence of KIT and that it is also functional *in vivo*, contrary to prior *ex vivo* findings.

Starting from E10.5, EMPs migrate into the fetal liver (Palis and Yoder, 2001), where they expand and differentiate. Our single cell transcriptomic analysis of E12.5 fetal liver demonstrated that KIT transcripts are enriched in liver erythroid progenitors of the BFU-E type (**Fig. 4**). Consistent with this expression pattern, we further found that the KIT null liver has a significant decrease of erythroid cells but concomitant increase of myeloid cells. Accordingly, KIT appears to regulate EMP fate decisions, whereby it is selectively required for transient-definitive erythropoiesis in the fetal liver downstream of EMP formation in the yolk sac (**Fig. 4**). Our study therefore provides new insights into the precise requirement of KIT for developmental hematopoiesis. Specifically our study refines previous work based on *Kit^W/W^* and *Kitl^Sl/Sl^* spontaneous mutants (Chui et al., 1978; Chui and Russell, 1974; Russel et al., 1968) and neutralizing anti-KIT antibodies (Ogawa et al., 1993), which showed an erythroid differentiation block in the fetal liver after E13.5 without identifying the specific progenitor population involved, because we have identified fetal liver erythroid differentiation from EMP-derived progenitors as the first event in which KIT acts to prevent severe anemia. Our observations agree with a recent study of mice lacking the KIT ligand KITL, which is also known as stem cell factor (SCF); specifically, this prior study reported a striking decrease in EMP-derived fetal liver erythroid cells at E12.5 (Azzoni et al., 2018). Our work also shows that KIT is dispensable for the formation of microglia (**Fig. 3**), which differentiate from yolk sac hematopoietic progenitors that arise in the yolk sac before the birth of the liver-colonizing EMPs (Ginhoux and Guilliams, 2016; Hoeffel and Ginhoux, 2018); these observations also agree with findings in mice lacking KITL (Azzoni et al., 2018).

Our transcriptomic profiling of the E12.5 fetal liver further demonstrated that *Kit* expression is limited to hematopoietic and erythroid-biased progenitors, with no detectable expression in ECs at this stage (**Fig. 4**). Lack of *Kit* expression in E12.5 liver was unexpected, because brain ECs express *Kit* at E12.5 (**Fig. 3**), and because *Kit* is abundantly expressed in adult hepatic sinusoidal ECs (Mansuroglu et al., 2009; see also Tabula Muris database, https://tabula-muris.ds.czbiohub.org/, Tabula Muris et al., 2018). Our findings therefore suggest that *Kit* expression in liver ECs is acquired later on during development, perhaps after the liver vasculature has specialized into sinusoids that connect the portal triads to the central veins (Strauss et al., 2017).

As our scRNAseq analysis corroborated a recent report suggesting that *Kit* is expressed in embryonic brain ECs during brain angiogenesis (Feng et al., 2020), we asked whether KIT promotes brain angiogenesis. However, embryonic brain vascularization in the well-established hindbrain angiogenesis model was not obviously affected by KIT loss (**Fig. 3**). This was unexpected, because KIT has previously been shown to promote the migration, proliferation and tube formation of cultured endothelial cells (Kim et al., 2011; Kim et al., 2019; Matsui et al., 2004), and because decreased KIT expression reduces angiogenesis in mouse models of ocular pathology (Kim et al., 2019) and tumor growth (Fang et al., 2012). The KIT requirement for angiogenesis under hypoxic and inflammatory pathological conditions was recently ascribed to the hypoxia-induced upregulation of KIT protein levels, which activates β-catenin pro-angiogenic signaling in ECs (Kim et al., 2019).

Although endothelial *Kit* expression would agree with two recent reports describing multilineage hematopoietic cells arising from brain ECs between E10.5 and E13.5 (Gama Sosa et al., 2020; Li et al., 2012), we did not detect any transcript for the hemogenic EC marker *Runx1* in the brain EC population at either E11.5 or E12.5 (**Fig. 2**). Lack of *Runx1* expression implies that endoHT does not occur in the brain, at least at this developmental stage. Future work should therefore examine whether hemogenic ECs exist in the brain at earlier developmental stages. Alternatively, hematopoietic progenitors might only be contained within developing brain vasculature, rather than arising from brain hemogenic endothelial cells.

The embryonic brain contained non-EC and non-hematopoietic cells that were positive for *Kit*, including neural progenitors and neuroblasts (**Fig. 2**). In agreement with *Kit* expression in neural progenitors, lineage tracing driven from the endogenous *Kit* promoter in E8.5 *Kit^CreERT2^;Rosa^tdTom^* embryos identified *Kit* lineage neural cells in the E12.5 hindbrain (Plein et al., 2018). *Kit* is therefore expressed by neural progenitors from around E8.5 onwards and maintained to at least E12.5. It would therefore be interesting to examine whether KIT contributes to neurogenesis in the central nervous system, in analogy to its role in the neural crest-derived neurons of the peripheral nervous system, whereby it was previously shown that KIT promotes the survival and neurite outgrowth of sensory neurons in the dorsal root ganglia (Hirata et al., 1993).

The most important and definite finding of our study is the requirement of KIT for transient-definitive fetal liver erythropoiesis downstream of EMP formation in the yolk sac, with a key role in regulation of erythroid versus myeloid fate (see working model, **Fig. 4j**). In agreement with our observation that KIT promotes erythroid at the expense of myeloid cell fate, gain-of-function mutations in the *Kit* coding sequence have been described to trigger clonal expansion of malignant pro-erythroblasts in murine erythroleukemia (Kosmider et al., 2005). Moreover, gain-of-function mutations have been associated with human adult and pediatric core binding factor acute myeloid leukemia (CBF-AML), for which KIT mutations are poor prognostic factors (Cairoli et al., 2006; Chen et al., 2018; Krauth et al., 2014). It should therefore be investigated whether *KIT* mutations also contribute to human erythroleukemia.

## Method

### Mouse strains

All animal procedures were performed in accordance with the institutional Animal Welfare Ethical Review Body (AWERB) and UK Home Office guidelines. To obtain mouse embryos of defined gestational age, mice were paired in the evening and the presence of a vaginal plug the following morning was defined as E0.5. Mice carrying the *Csf1r-iCre* transgene (MGI:4429470; Deng et al., 2010) were mated to mice carrying the *Cre* recombination reporters *Rosa^Yfp^* (MGI:2449038; Srinivas et al., 2001) or *Rosa^tdTom^* (MGI:3809524; Madisen et al., 2010). Embryos lacking *Kit* expression were generated by mating *Kit^CreERT2^* mice (MGI:5543260; Klein et al., 2013) or by mating *Kit^CreERT2^* mice carrying *Rosa^tdTom^* and *Csf1r-iCre* to obtain *Csf1r-iCre;Rosa^tdTom^;Kit*^−/−^ embryos. All mouse strains were maintained on a mixed background (C57Bl6/J;129/Sv, C57Bl6/J;FVB or C57Bl6/J;129/Sv;FVB).

### Wholemount tissue staining

Samples were fixed in 4% formaldehyde in PBS and processed as wholemounts for fluorescent staining as described previously for wholemount hindbrains (Fantin et al., 2013b). We used the following antibodies and dilutions: rat anti-CDH5 (1:200; 555289, BD Pharmingen), rat anti-F4/80 (1:500; MCA497R, lot 1605, Serotec), rat anti-KIT (1:500; 553353, lot 30259, BD Pharmingen), rabbit anti-RFP (1:1,000; PM005, lot 045, MBL), goat anti-VEGFR2 (1:200; AF644, lot COA0417021, R&D Systems). Secondary antibodies used included donkey fluorophore-conjugated FAB fragments of anti-goat, -rabbit or -rat IgG (Jackson ImmunoResearch). Biotinylated IB4 (1:200; L2140, lot 085M4032V, Sigma) followed by Alexa-conjugated streptavidin (ThermoFisher) was used to detect brain ECs (Fantin et al., 2013a). Images were acquired with a LSM710 (Zeiss) or an A1 (Nikon) laser scanning confocal microscope and processed using LSM image browser (Zeiss), Fiji (NIH Bethesda(Schindelin et al., 2012) and Photoshop CS4 (Adobe Inc.) software. Three-dimensional reconstructions including surface rendering and the generation of virtual slices for lateral views of high-resolution confocal *z*-stacks was performed using Imaris (Bitplane). *Z*-stack projections of confocal images are shown unless indicated otherwise in the figure legends.

### Flow cytometry

Tissues were mechanically and enzymatically homogenized in RPMI1640 with 2.5% fetal bovine serum (ThermoFisher), 100 μg/ml collagenase/dispase (Roche), 50 μg/ml DNase (Qiagen) and 100 μg/ml heparin (Sigma), incubated for 5 min with 0.5 mg/ml rat Fc block (Becton Dickinson) and labelled with a combination of BV585-, PerCp/Cy5.5-, PE-Dazzle 594-or FITC-conjugated rat anti-CD45 (clone 30-F11, cat 103108, lot B246762) or FITC-conjugated rat anti-CD41 (clone MWReg30, cat 133903, lot B201955), PerCp-Cy5.5-or APC-conjugated rat anti-KIT (clone 2B8, cat 105812, lot B217855), FITC-conjugated rat anti-TER-119 (clone TER-119, cat 116206) and PerCp-Cy5.5-conjugated rat anti-CD11b (clone M1/70, cat 101227) (all BioLegend). Appropriate fluorescence gating parameters were established with unstained or *Csf1r-iCre*-negative tissues and fluorescence-minus-one (FMO) staining. In all experiments, doublets were eliminated using pulse geometry gates (FSC-H versus FSC-A and SSC-H versus SSC-A), whereas dead cells were removed using SYTOX Blue (Life Technologies) or LIVE/DEAD Fixable Violet (Life Technologies). Single-cell suspensions were analysed using the BD LSRFortessa X-20 cell analyser or sorted using the BD Influx cell sorter (BD Biosciences); FlowJo software (FlowJo LLC) was used for subsequent analyses.

### Single cell RNA sequencing (scRNA-seq) analysis

Raw data for the E11.5 and E12.5 midbrain scRNA-seq datasets were obtained from GSE76381 (NCBI GEO, La Manno et al., 2016). Analyses were performed with RStudio 3.6.3. The raw gene expression matrices (UMI counts per gene per cell) were filtered, normalised and clustered using the R package Seurat 3.2.0. Cells containing less than 200 feature counts and genes detected in less than 3 cells were removed, obtaining 406 cells for the E11.5 dataset and 350 for the E12.5 dataset. Downstream analysis included data normalization (“LogNormalize” method and scale factor of 10,000) and variable gene detection (“vst” selection method, returning 2,000 features per dataset). The principal components analysis (PCA) was performed on variable genes, and the optimal number of principal components, PCs, for each sample was chosen using the elbow plot (15 for both datasets). Louvain graph-based clustering was performed using the selected PCs and a resolution of 2.2 for the E11.5 dataset and 5.5 for the E12.5. Dimensionality reduction methods included UMAP and cluster cell identity was assigned by manual annotation using known marker genes. The E12.5 liver scRNA-seq dataset was generated from C57BL/6J fetal liver (manuscript in preparation). Briefly, a single cell suspension from one E12.5 liver, obtained by enzymatic digestion and mechanical dissociation was purified from debris, dead cells and doublets through fluorescence activated cell sorting (FACS). To maximise the yield in droplet encapsulation of single cells, two separate technical replicates of the sample were independently processed. Next, the library was prepared using the Chromium Single-Cell Controller (10X Genomics, Pleasanton, CA) and sequenced at a depth of 50k reads/cell on a NovaSeq 6000 (Illumina). Raw files were processed with Cell Ranger 2.1.1 and reads mapped to the mm10 genome and counted with GRCm38.92 annotation. Using Seurat, cells containing feature counts less than 200 feature counts and genes detected in less than 3 cells were removed, obtaining 5,480 and 5,057 cells, respectively, for the two technical replicates, which were then merged into a single sample for subsequent analysis. Normalization, PC selection (15) and Louvain graph-based clustering were applied as described for the midbrain datasets with a resolution of 0.6. Cluster cell identity was assigned by manual annotation using known marker genes, including those shown in Figure 4.

### Statistical analysis

Tissues for analysis were allocated to experimental groups according to genotype, gestational age, organ or cell type, rather than being randomized. To ensure the unbiased interpretation of results, the genotype and gestational age were disclosed only after data collection was complete, but the investigators knew the sample origin (i.e. organ or cell type). No statistical methods were used to predetermine sample size. For hindbrains experiments, the number of F4/80^+^ microglia and the tdTomato^+^ and IB4^+^ volumes were determined from confocal *z*-stacks of four randomly chosen 0.25 mm^2^ regions on the lateral side of each hindbrain. The *z*-stacks were surface rendered with Imaris (Bitplane) to obtain the F4/80^+^, tdTom^+^ and IB4^+^ volumes, and the F4/80^+^ volume was then subtracted from both the IB4^+^ and tdTom^+^ total volumes to obtain the IB4^+^ EC and tdTom^+^ EC volumes before calculating the ratio of tdTom^+^ to IB4^+^ EC volume. The same confocal *z*-stacks were analysed with Angiotool (Zudaire et al., 2011) to determine the number of vascular intersections and with Fiji (Schindelin et al., 2012) to count the number of F4/80^+^ microglia. All counts obtained from one hindbrain were averaged to yield the value for that hindbrain. For all experiments, the error bars represent the standard deviation of the mean (for details, see legends). Comparison of medians against means justified the use of a parametric test; to determine whether two datasets were significantly different, we therefore calculated *P* values with a two-tailed unpaired Student’s *t*-test; *P*< 0.05 was considered significant. Statistical analyses were performed with Excel 12.2.6 (Microsoft Office) or Prism 7 (GraphPad Software).

## Author Contributions

AF and CR contributed to conception and design of the study and co-wrote the manuscript. AF, AP and LD performed mouse experiments. AF and CT analyzed data. EV and EC performed bioinformatic analyses. All authors read and approved the submitted manuscript.

## Acknowledgments

We thank the staff of the Biological Resources, FACS and Imaging Facilities at the UCL Institute of Ophthalmology and the FACS and Genomics facilities at the UCL Institute of Child Health for technical help. We thank Alison Domingues for helpful comments on the manuscript.

## Data Availability

Publicly available datasets were analyzed in this study. Midbrain scRNAseq data can be found here: [https://www.ncbi.nlm.nih.gov/geo/, GSE76381]. Fetal liver scRNAseq data can be found here: [https://www.ncbi.nlm.nih.gov/geo/, available upon publication]. Further inquiries can be directed to the corresponding authors.

## Contribution to the Field Statement

A combination of mouse genetics and transcriptomic analyses revealed that the receptor tyrosine kinase KIT is required for transient definitive erythropoiesis in the fetal liver, but dispensable for the function of yolk sac hemogenic endothelium and its progeny, the yolk sac-born erythromyeloid progenitors (EMPs) that give rise to the progenitors of transient definitive erythrocytes in the fetal liver. We further found that KIT is expressed in embryonic brain endothelial cells but is dispensable for brain vascularization. KIT expression in hemato-endothelial cells is therefore not always indicative of a functional requirement. Instead, hemato-endothelial KIT expression is selectively required for transient definite erythropoiesis in the fetal liver (see graphical abstract).

